# Kupffer cell and recruited macrophage heterogeneity orchestrate granuloma maturation and hepatic immunity in visceral leishmaniasis

**DOI:** 10.1101/2024.07.09.602717

**Authors:** Gabriela Pessenda, Tiago R. Ferreira, Andrea Paun, Juraj Kabat, Eduardo P. Amaral, Olena Kamenyeva, Pedro Henrique Gazzinelli-Guimaraes, Shehan R. Perera, Sundar Ganesan, Sang Hun Lee, David L. Sacks

**Affiliations:** Laboratory of Parasitic Diseases, National Institute of Allergy and Infectious Diseases, National Institutes of Health, Bethesda, MD 20892, USA; Biological Imaging Section, Research Technology Branch, National Institute of Allergy and Infectious Diseases, National Institutes of Health, Bethesda, MD 20892, USA; Department of Electrical and Computer Engineering, The Ohio State University, Columbus, OH 43201, USA

**Keywords:** Kupffer cells, visceral leishmaniasis, monocyte-derived macrophages, granuloma, ferroptosis, chronic infection, parasitic diseases, cell death, liver

## Abstract

In murine models of visceral leishmaniasis (VL), parasitization of resident Kupffer cells (resKCs) is responsible for early growth of *Leishmania infantum* in the liver, which leads to granuloma formation and eventual parasite control. We employed the chronic VL model, and revealed an open niche established by KCs death and their migration outside of the sinusoids, resulting in their gradual replacement by monocyte-derived KCs (moKCs). While early granulomas were composed of resKCs, late granulomas were found outside of the sinusoids and contained resKC-derived macrophages, and monocyte-derived macrophages (momacs). ResKCs and moKCs within the sinusoids were identified as homeostatic/regulatory cells, while resKC-derived macrophages and momacs within late granulomas were pro-inflammatory. Despite the infection being largely confined to the resKC-derived macrophages, in the absence of monocyte recruitment, parasite control was strongly compromised. Macrophage heterogeneity, involving migration and reprogramming of resKCs, along with recruitment of monocyte-derived cells, is a hallmark of granuloma maturation and hepatic immunity in VL.

## Main

Kupffer cells (resKCs) are the embryonic-derived, tissue-resident macrophages (TRMs) that lie within the liver sinusoids and remain non-migratory during homeostatic conditions^1–5^. In some inflammatory settings or when deliberately targeted for depletion, they can be replaced by monocyte-derived KCs (moKCs)^6–17^. Granulomas are organized aggregates of macrophages and other immune cells that arise in response to infectious or non-infectious agents^18^. Experimental visceral leishmaniasis (VL) caused by *Leishmania infantum/donovani* is a well characterized chronic infection model to study granuloma dynamics and function in the liver^19^. Infected KCs initiate granuloma formation during VL, involving KC aggregation and fusion, along with recruitment of innate and adaptive immune cells^20,21^. The possible heterogeneity of granuloma macrophages regarding their ontogeny and function has not been addressed.

Kupffer cells in naïve mice are characterized as F4/80^hi^CD11b^int^CD64^+^CLEC4F^+^TIM-4^+^ (Extended Data Fig.1a). While CLEC4F is a KC specific marker, TIM-4 expression on moKCs is delayed, and distinguishes KCs ontogeny ^6–8,12,14,22,23^. At forty-two days post *L. infantum* infection (42 d.p.i.), we observed 4 subsets of F4/80^hi^CD11b^int^CD64^+^ liver macrophages in C57BL/6 mice. While CLEC4F^+^TIM-4^+^resKCs frequency was reduced, CLEC4F^+^TIM-4^-^moKCs, and CLEC4F^-^TIM-4^+^ and CLEC4F^-^TIM-4^-^ macrophages were all increased compared to naïve mice (Extended Data Fig.1b-c). To investigate these subsets *in situ* at various stages of *L. infantum-* induced liver inflammation, we used confocal microscopy on liver sections at different time points (Fig. 1a). Most F4/80^+^ macrophages in naïve and 19 d.p.i. mice, defined here as “early infection”, were CLEC4F^+^TIM-4^+^resKCs. These cells were reduced in frequency and number in 42 d.p.i. livers (“late infection”), coinciding with the peak of CLEC4F^-^TIM-4^+^ and CLEC4F^-^TIM-4^-^ populations. CLEC4F^+^TIM-4^-^moKCs continued to accumulate during the “resolving phase” (72 d.p.i.) (Fig. 1a-b). Similar to the observation in whole liver tissue, the inset analysis of the cellularity within the granulomas revealed that CLEC4F^+^TIM-4^+^resKCs were the principal subset found within early forming granulomas at 19 d.p.i., but were greatly reduced in 42-day granulomas, which contained mainly CLEC4F^-^TIM-4^+^ and CLEC4F^-^TIM-4^-^ cells. In addition, seventy-two-day granulomas, while few in number, contained roughly equivalent ratios of all four F4/80^+^ subsets (Fig. 1c-d). Overall, resKCs were significantly reduced during late VL in favor of F4/80^+^ macrophages that did not express CLEC4F and/or TIM-4. While early infection granulomas contained mainly resKCs, later infection resulted in granulomas containing heterogeneous macrophage populations.

**Figure 1:**
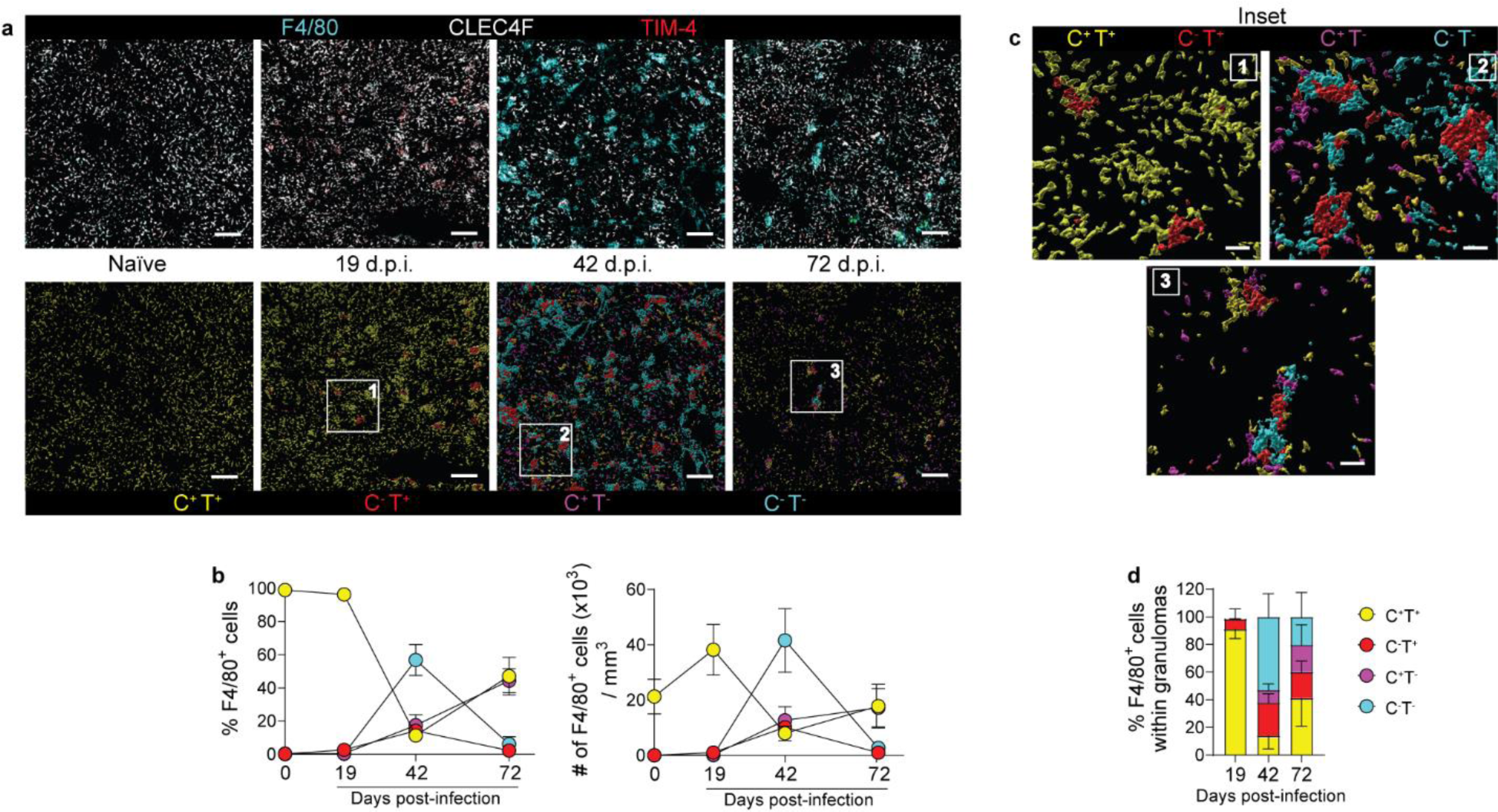
Heterogeneity of F4/80^+^ macrophages during VL defined by the differential expression of CLEC4F and TIM-4. **a**, Representative confocal microscopy images showing WT naïve, 19-, 42-, and 72-day infected livers stained with anti-F4/80 (cyan), anti-CLEC4F (white), and anti-TIM-4 (red), and surface rendered F4/80^+^ cells classified according to their CLEC4F and TIM-4 expression: CLEC4F^+^TIM-4^+^ (yellow), CLEC4F^-^TIM-4^+^ (red), CLEC4F^+^TIM-4^-^ (magenta), CLEC4F^-^TIM-4^-^ (cyan). Scale bars, 200μm. **b**, Frequency and number of F4/80^+^ cells classified according to their CLEC4F and TIM-4 expression, in naïve and infected WT mice at different days post-infection. Data pooled from 2 independent experiments (n=6 for naïve and n=7 for infected mice). **c**, Inset shows granulomas identified in (a) and defined as clusters of F4/80^+^ cells with volumes above 1.03×10^4^ μm^3^. Scale bars, 40μm. **d**, Proportion of CLEC4F^+^ and/or TIM-4^+^ cells in F4/80^+^ granulomas at different times post-infection. Data pooled from 2-4 independent experiments (n=7 for 19- and 72 d.p.i. and n=15 for 42 d.p.i.). Values in **b** and **d** represent the mean ± SD.

To confirm the origin of the macrophage subsets identified in the liver at 42 d.p.i., we generated congenically paired, parabiotic mice, which shared a chimeric blood supply. In naïve mice, almost all the F4/80^hi^CD11b^int^CD64^+^ macrophages bore the congenic marker of the host parabiont, corroborating their origin as TRMs. At 42 d.p.i., 10.5±7.8% of the cells originated from the congenic partner (Fig. 2a-b). When further classifying CD45.2 cells within infected livers of the CD45.1 parabiont, CLEC4F^+/-^TIM-4^-^ macrophages and monocytes contained cells from the congenic partner, while CLEC4F^+/-^ TIM-4^+^ macrophages were still mainly from the host parabiont (Fig. 2c), providing strong evidence that the absence of TIM-4 expression identifies macrophages of monocytic origin during VL. We also used *Ccr2*^-/-^ mice, in which *Ccr2^+^* monocyte recruitment to the liver was greatly reduced in both naïve (Fig. 2d) and infected mice (Fig. 2e). By confocal microscopy, although infected WT and *Ccr2*^-/-^ mice showed similar F4/80^+^ macrophage numbers (Fig. 2f), CLEC4F^+/-^TIM-4^-^ subsets frequencies were drastically reduced, and CLEC4F^+/-^TIM-4^+^ were proportionately increased compared to WT mice (Fig. 2g). These changes reflected overall differences in macrophage heterogeneity, with infected WT mice showing all four F4/80^+^ subsets, while infected *Ccr2*^-/-^ mice showed mostly TIM-4^+^ macrophages and granulomas (Fig. 2h-i and Extended Data Fig.2a). The presence of CLEC4F^-^TIM-4^+^ macrophages in granulomas of the *Ccr2^-/-^* mice, and parabiosis that showed these cells were host derived, suggested they could be CLEC4F^+^TIM-4^+^resKCs that downregulated CLEC4F expression inside granulomas. By breeding *Clec4f-Cre-tdTomato* mice with *RCL-ZsGreen* mice, which have a *loxP*-flanked STOP cassette preventing transcription of CAG promoter-driven *ZsGreen,* we were able to distinguish active CLEC4F expression marked by tdTomato from prior expression indicated by ZsGreen positivity. In naïve mice, resKCs were tdTomato^+^ZsGreen^+^ (Fig. 2j). At 42 d.p.i, tdTomato^+^ZsGreen^+^KCs were found as individual cells inside the sinusoids, while tdTomato^-^ZsGreen^+^ cells were found in clusters of cells outside the sinusoids (Fig. 2k-l), confirming that resKCs downregulate CLEC4F inside granulomas. Because they no longer expressed the KC specific marker, we referred to the CLEC4F^-^TIM-4^+^ macrophages as resKC-derived macrophages. Taken together, infected WT livers showed a heterogeneous macrophage population consisting of CLEC4F^+^TIM-4^+^resKCs, CLEC4F^-^TIM-4^+^resKC-derived macrophages, CLEC4F^+^TIM-4^-^moKCs and CLEC4F^-^TIM-4^-^momacs, whereas infected *Ccr2*^-/-^ livers contained only resKCs and resKC-derived macrophages.

**Figure 2:**
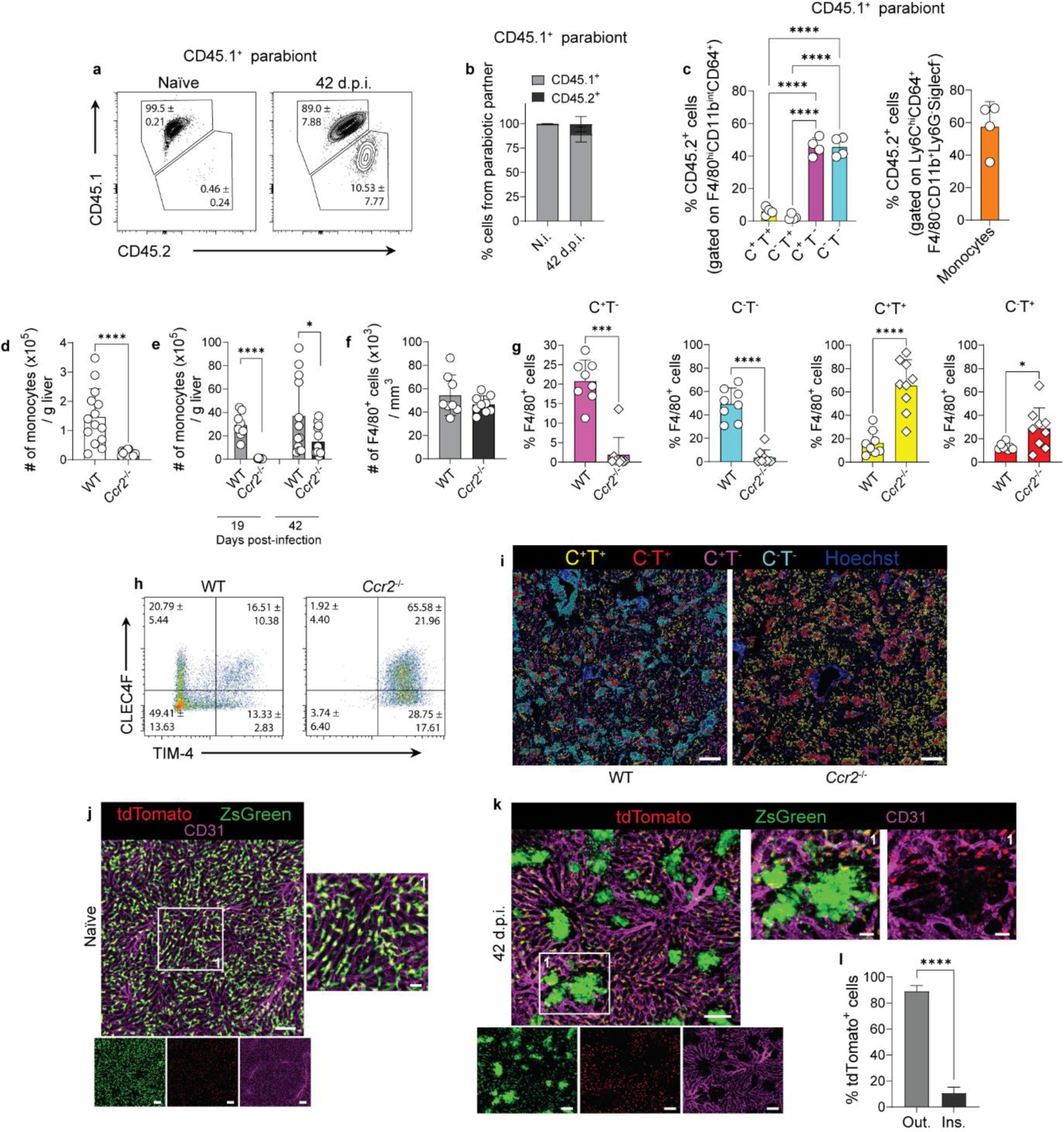
TIM-4^-^ macrophages are monocyte-derived at 42 d.p.i. **a**, Representative contour plots showing the percentages of chimerism in F4/80^+^CD11b^int^CD64^+^ cells of naïve and 42 days infected CD45.1^+^ parabiont. **b**, Percentages of chimerism in F4/80^+^CD11b^int^CD64^+^ cells of uninfected and infected CD45.1^+^ parabiotic partner at 42 d.p.i. **c**, Frequency of CD45.2^+^ cells in infected CD45.1^+^ parabiont and gated on F4/80^+^CD11b^int^CD64^+^CLEC4F^+/-^TIM-4^+/-^ macrophages or on Ly6C^+^CD11b^+^Ly6G^-^ SiglecF^-^ monocytes. Data pooled from two independent experiments (n=3 for uninfected and n=4 for infected pairs). **d-e**, Number of monocytes in the liver of naïve (**d**) and 19- and 42-day infected WT and *Ccr2*^-/-^ mice (**e**), obtained by flow cytometry. Data pooled from 5 independent experiments (naïve, n=14 for WT and n=17 for *Ccr2*^-/-^; 19 d.p.i., n=8 for WT and n=9 for *Ccr2*^-/-^; 42 d.p.i., n=12 for WT and n=14 for *Ccr2*^-/-^). **f**, Number of F4/80^+^ cells in WT and *Ccr2*^-/-^ at 42 d.p.i., quantified from confocal microscopy images. **g**, Frequency of macrophage subsets based on CLEC4F and TIM-4 expression in WT and *Ccr2*^-/-^ mice at 42 d.p.i., quantified from confocal microscopy images. **h**, Representative dot plots from confocal microscopy images showing the frequency of macrophage subsets in infected WT and *Ccr2*^-/-^ mice at 42 d.p.i. Numbers indicate mean ± SD percentage of cells in the gate. **i**, Representative rendered confocal microscopy images of WT and *Ccr2*^-/-^ 42 d.p.i. livers showing CLEC4F^+^TIM-4^+^ (yellow) and CLEC4F^-^TIM-4^+^ (red) macrophages, CLEC4F^+^TIM-4^-^ (magenta) moKCs and CLEC4F^-^TIM-4^-^ (cyan) momacs. Scale bars, 200μm. Data pooled from 2 independent experiments (n=8 for WT and n=9 for *Ccr2*^-/-^). **j-k**, Representative live images of naïve (**j**) and 42-day infected (**k**) *Clec4f-Cre-ZsGreen* mice showing Clec4f active expression (red), Clec4f previous expression (green), and sinusoids (magenta). Scale bars, 100μm and 30μm (insets). **l**, Frequency of tdTomato^+^ cells identified inside and outside granulomas, at 42 d.p.i. Data pooled from 2 independent experiments (n=4). Values in **b**-**g** and **l** show mean ± SD. *, p < 0.05, ***, p < 0.001, ****, p < 0.0001.

It has been suggested that spatial redistribution of KCs along the sinusoidal network, as has been described in *L. donovani*^24^, *L. monocytogenes*^9^, and *Mycobacterium bovis* BCG infected livers, can result in an open KC niche^25^. Live imaging of naïve *Clec4f-Cre-ZsGreen* mice showed that tdTomato^+^ZsGreen^+^KCs were inside or in close contact with the sinusoids (Fig. 2j). Using additional intravenous F4/80 staining in 42-day infected mice, while the majority of F4/80^+^ macrophages inside the sinusoids were tdTomato^+^ZsGreen^+^ and tdTomato^-^ZsGreen^-^, a few were tdTomato^-^ZsGreen^+^ (Fig. 3a-b). As observed in Fig. 2k and Fig. 3c, infection resulted in an intense sinusoidal remodeling, with granulomas outside of the sinusoids containing mostly clusters of tdTomato^-^ ZsGreen^+^ (Fig. 3c-d). When detected inside granulomas, tdTomato^+^ZsGreen^+^KCs were closer to the sinusoids (Fig. 3c). Intravascular staining allowed the discrimination between tissue and blood-borne cells^26^. Most ZsGreen^+^ cells inside granulomas did not stain for F4/80, while the few that we detect partially inside or close to the sinusoids were stained, suggesting their recent migration (Fig. 3c). Most resKCs that migrated outside of the sinusoids and were at the core of the granulomas were no longer in contact with LSECs and converted into tdTomato^-^ZsGreen^+^resKC-derived macrophages. Considering that KC interaction with hepatocytes, hepatic stellate cells (HSCs), and LSECs in the liver is important for their survival^6^, it is possible that lack of contact between KCs and LSECs was responsible for their loss of identity, evidenced by CLEC4F downregulation. Loss of KC identity has been described in studies targeting genes involved in its maintenance, including *Alk1* and *Smad4*^27^, and during liver fibrosis^28^, which resulted in increased frequencies of CLEC4F^-^TIM-4^+^KCs.

**Figure 3:**
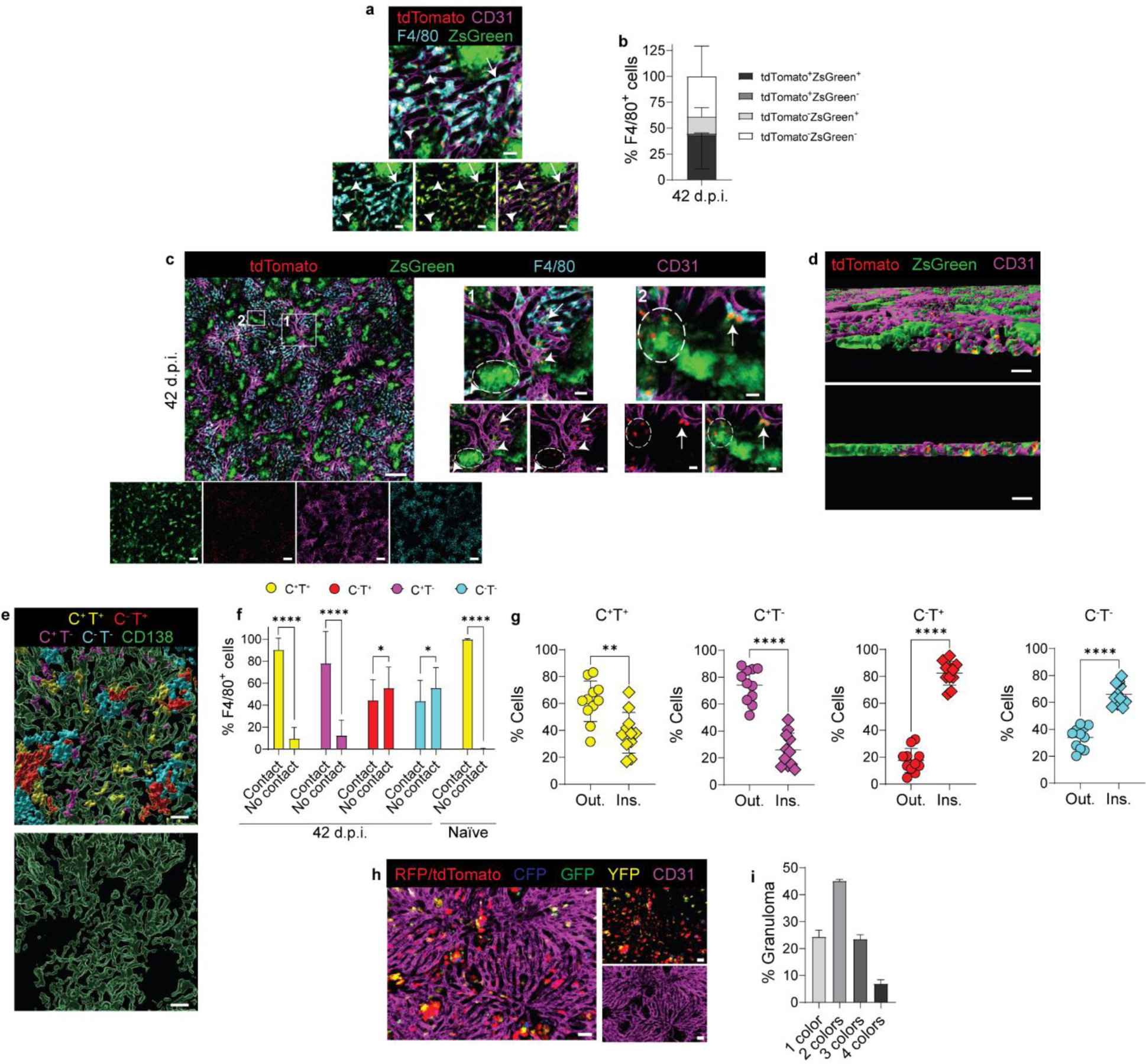
Late granulomas form outside of the sinusoids and contain polyclonal clusters of macrophages derived from res-KCs. **a**, Representative live images from a 42-day infected *Clec4f-Cre-ZsGreen* mouse showing Clec4f active expression (red), Clec4f previous expression (green), F4/80 (cyan), and sinusoids (magenta). Arrow shows a F4/80^+^tdTomato^-^ZsGreen^+^ cell, and arrowheads show F4/80^+^tdTomato^-^ZsGreen^-^ cells. Scale bars, 30μm. **b**, Frequency of F4/80^+^ cells inside the sinusoids according to tdTomato and ZsGreen expression, in 42 d.p.i. mice. **c**, Representative images of *Clec4f-Cre-ZsGreen* mice at 42 d.p.i., showing Clec4f active expression (red), Clec4f previous expression (green), F4/80 (cyan), and sinusoids (magenta). Scale bars, 200μm. Inset #1, dotted line shows clustered tdTomato^-^ZsGreen^+^ cells in granulomas outside the sinusoids. Arrowheads show one F4/80^+^ and one F4/80^-^ tdTomato^+^ZsGreen^+^ cell outside the sinusoids but in close contact with LSECs. Arrow points to 3 F4/80^+^tdTomato^+^ZsGreen^+^ cells outside the sinusoids. Scale bars, 30 μm. Inset #2, dotted lines show a tdTomato^-^ZsGreen^+^ cluster with 3 tdTomato^+^ZsGreen^+^ closer to the LSECs. Arrow points to 2 tdTomato^+^ZsGreen^+^ partially inside the sinusoids and partially stained with F4/80. Scale bars, 15 μm. Data pooled from 2 independent experiments (n=4) **d**, Representative rendered images showing tdTomato (red) and ZsGreen (green) expression inside and outside the sinusoids (magenta), in 42-day infected *Clec4f-Cre-ZsGreen* mouse. Scale bars, 80 μm. **e**, Representative immunofluorescence image of a 42-day infected WT liver showing the spatial distribution of CLEC4F^+^TIM-4^+^ resKCs (yellow), CLEC4F^-^TIM-4^+^ resKC-derived macrophages (red), CLEC4F^+^TIM-4^-^ moKCs (magenta), CLEC4F^-^TIM-4^-^ momacs (cyan), and LSECs (green). Scale bars, 30 μm. **f**, Bar graphs showing the localization of each macrophage subset according to their interaction with LSECs, in 42 d.p.i. and naïve mice. Data pooled from 2 independent experiments (n=4 for naïve and n=8 for 42 d.p.i.). (Quantification of 4 regions of interest (ROIs) from each infected mouse, n=32 ROIs, and 2-3 ROIs from each uninfected mouse, n=11 ROIs). **g**, Scatter plots from immunofluorescence images showing the frequency of each F4/80^+^ population based on their distribution outside or inside granulomas at 42 d.p.i. Data pooled from 3 independent experiments (n=11). **h,** Intravital microscopy images representing a 42-day infected liver from *Clec4f-Cre-Confetti* mouse stained with anti-CD31 (magenta) and showing tdTomato (red-nucleus), RFP (red-cytoplasm), YFP (yellow-cytoplasm), GFP (green-nucleus), and CFP (blue-membrane) KCs. Scale bars, 50 μm. **i,** Composition of macrophage granulomas based on KC and KC-derived macrophages colors. Data pooled from live imaging carried out in 2 independent experiments (n=4 and quantification of 284 granulomas). Values from **b**, **f-g** and **i** represent mean ± standard deviation. *, p < 0.05, **, p < 0.01, ****, p < 0.0001.

While live imaging of *Clec4f-Cre-ZsGreen* infected livers allowed us to distinguish resKCs from resKC-derived macrophages, it did not discriminate resKCs from moKCs, which are both CLEC4F^+^, or CLEC4F^-^TIM-4^-^momacs within granulomas. By confocal microscopy, we showed that CLEC4F^+^TIM-4^+^resKCs and CLEC4F^+^TIM-4^-^moKCs maintained their association with LSECs and were found mostly inside the sinusoids, while almost 60% of the CLEC4F^-^TIM-4^+^resKC-derived macrophages and CLEC4F^-^TIM-4^-^momacs did not have contact with LSECs and were outside of the sinusoids (Fig. 3e-f). Consistently, most resKCs and moKCs were outside of granulomas and resKC-derived macrophages and momacs were inside (Fig. 3g). To address the clonality of the KCs/resKC-derived macrophages within the granulomas, we bred *Clec4f-Cre-tdTomato* mice with *R26R-Confetti* mice, in which Cre recombinase causes permanent expression of one of four possible fluorescent proteins, allowing us to distinguish clonal lineages. At 42 d.p.i., approximately 70% of the granulomas contained 2 or more colors of resKC-derived macrophages (Fig. 3h-i), providing direct evidence that most granulomas were constituted by KCs that actively migrated to form polyclonal clusters and were not solely aggregates of self-proliferating cells.

Replacement of resKCs by moKCs may also occur after KC niche availability resulting from cell death^8,9,15,17,29^. Murine models of *Listeria monocytogenes*^9^ infection and viral hepatitis^11^ induce a massive loss of resKCs and rapid replacement by moKCs. In our chronic VL model, the replacement of the resKCs was more gradual, with CLEC4F^+^TIM-4^-^moKCs occupying the sinusoidal space and coexisting with the remaining resKCs at 42 d.p.i. To evaluate the possible roles of cell death in our VL model, we used mice deficient in MLKL, a terminal protein in the necroptotic cell death program, and mice deficient in Caspase 1, involved in the lytic, pyroptotic cell death pathway^30^. No differences in resKC or moKC frequencies were observed in *Mlkl^-/-^* (Extended Data Fig.3a) or *Casp1^-/-^* livers at 42 d.p.i. (Extended Data Fig.3b). Using cleaved caspase 3 staining to detect apoptotic cells *in situ*, we observed increased numbers of clv-casp3^+^ in all F4/80^+^ subsets and in F4/80^-^ cells at 42 d.p.i., except resKCs (Extended Data Fig.3c). While the majority of clv-casp3^+^ cells were F4/80^-^, the frequency of apoptotic cells among the resKCs was reduced during infection but increased in all other F4/80^+^ macrophages (Extended Data Fig.3d-e). Ferroptosis is an iron-dependent oxidative cell death involving peroxidation of lipids resulting in biological membrane damage^31^. It has been implicated in various hepatic diseases, including malaria^29,32^. By flow cytometry, we found evidence of ferroptotic cell death in both TIM-4^+^ and TIM-4^-^ macrophages at 42 d.p.i. (Extended Data Fig.4a). Glutathione peroxidase 4 (GPX4) mediates the reduction of phospholipid hydroperoxides by utilizing glutathione (GSH) as a co-factor of GPX4^31^, and lower levels of GPX4 and/or GSH are commonly associated with ferroptosis^33–36^. We found that GSH levels were reduced at 42 d.p.i. compared to uninfected controls (Extended Data Fig.4b). BACH1 is known as a pro-oxidant factor by repressing NRF2, a master regulator of host antioxidant responses^37,38^, and its deficiency has been shown reduce oxidative stress-mediated ferroptosis^38,39^. TIM-4^+^ and TIM-4^-^ cells from 42d.p.i. *Bach1*^-/-^ mice showed reduced levels of lipid peroxidation (Extended Data Fig.4c). ResKCs and total F4/80^+^ macrophages were increased in infected *Bach1*^-/-^ mice, while the frequencies of moKCs and momacs were reduced, with no changes in their absolute numbers (Extended Data Fig.4d-h). Collectively, apoptosis was increased during VL in all F4/80^+^ and F4/80^-^ cells except resKCs, while ferroptosis regulated resKCs and total F4/80^+^ macrophage numbers in infected livers. Taken together, our data suggest that ferroptosis of resKC and their migration outside the sinusoids to form polyclonal granulomas contribute to an open sinusoidal niche and their partial replacement by CLEC4F^+^TIM-4^-^moKCs that repopulate the sinusoids.

To further investigate molecular program and the heterogeneity of the cells identified by confocal microscopy and flow cytometry, we sorted F4/80^hi^CD11b^int^CD64^+^ macrophages and performed single-cell RNA sequencing, obtaining transcriptomes on a total of 7,673 cells, of which 7,352 were macrophages. We used the dataset from Remmerie *et al.*^14^ on the identification and nomenclature of macrophages in the liver as a reference for mapping our single-cell data onto the UMAP structure. UMAP projection and clustering resulted in 3 clusters identified in naïve and 6 clusters in infected mice (Extended Data Fig.5a). Re-clustering and manual annotation resulted in two distinct clusters of Transitioning monocytes in infected mice (Fig. 4a). The conserved transcriptomic signature of KCs described by Guilliams et al.^22^ confirmed that cells from ResKCs, MoKCs, and Transitioning monocytes1 clusters expressed KC signature genes (Fig. 4b). In naïve mice, the ResKCs cluster contained exclusively CLEC4F^+^TIM-4*^+^* expressing-resKCs that did not express monocytic markers. In infected mice, in addition to CLEC4F^+^TIM-4*^+^*resKCs, this cluster also contained macrophages of a monocyte origin, given their expression of *Ccr2*/*Cxc3cr1* and/or lack of *Clec4f*/*Timd4* expression, suggesting these were monocyte-derived KCs transcriptionally similar to resKCs but still lacking the expression of resKC specific markers CLEC4F/TIM-4 (Fig. 4c-d and Extended Data Fig. 5b). Proliferating macrophages were increased in infected mice, and showed a gene expression profile more similar to ResKCs/MoKCs clusters, suggesting the majority of proliferating cells belonged to the KC compartment (Fig.4a,d-e and Extended Data Fig. 5b). Remmerie *et al.*^14^ identified the Mac1 cluster as pre-moKCs *en route* to becoming KCs because of their *Clec1b* expression, a marker for moKCs expressed earlier than *CLEC4F*^8^. We did not detect either *Clec1b* or KC signature genes in cells from this cluster (Fig. 4b,d). We cannot, however, exclude the possibility that they would eventually differentiate into KCs in infected mice, since they expressed several KC-associated transcription factors (Extended Data Fig. 5c). The Mac2 cluster was previously identified as hepatic lipid-associated macrophages (hep-LAMS)^14^, and we detected higher expression of *Spp1*, *Cd9* and *Trem2*, as expected for these macrophages (Fig. 4f). Finally, the term Transitioning monocytes was used for cells that showed features of both monocytes and macrophages^14^. The fact that cells from Transitioning monocytes1 cluster expressed KC signature genes and several KC transcription factors (Fig. 4b and Extended data Fig. 5c) may also indicate that these macrophages were committed to a KC fate. Of note, we could not identify a defined cluster of CLEC4F^-^TIM-4^+^ cells, which were well represented in the 42 d.p.i. livers by confocal microscopy (Fig. 1). Their absence may be explained by the recovery bias of the cells submitted for *ex vivo* analysis, which as recently discussed^40^, can result from liver disaggregation and digestion, resulting in under-representation of KCs and resKC-derived macrophages vs monocyte-derived populations. It is also possible that these cells were present within the Transitioning monocytes1 cluster, given their similar KC signature and *Nos2* expression.

**Figure 4:**
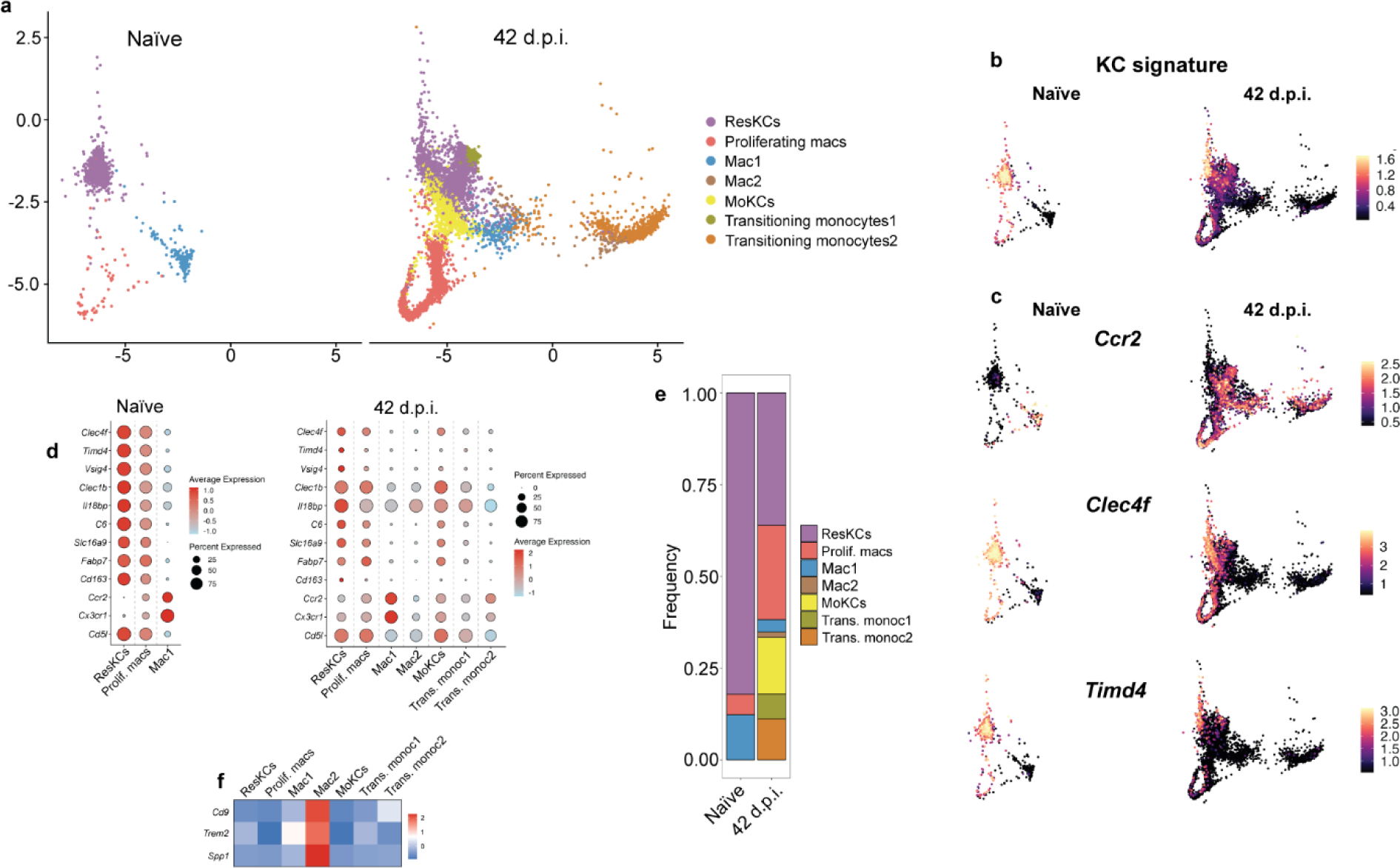
Macrophage heterogeneity revealed by single-cell RNA sequencing. **a,** UMAP plot of scRNA-seq data of sorted live, single, CD45.2^+^F4/80^+^CD11b^int^CD64^+^ cells from livers of uninfected and 42 d.p.i. mice, showing 3 clusters for naïve and 7 clusters for infected mice. **b,** Annotated UMAP plot showing the single-cell expression of conserved KC signature genes described by Guilliams et al.^22^ **c,** Single-cell expression of *Ccr2*, *Clec4f* and *Timd4* in CD45.2^+^F4/80^+^CD11b^int^CD64^+^ cells from naïve and 42 d.p.i. mice **d,** Average gene expression and percent of cells in which they are expressed within each of the identified clusters in naïve and 42 day-infected CD45.2^+^F4/80^+^CD11b^int^CD64^+^ cells. **e,** Frequency of the different subsets identified by scRNA-seq in naïve and 42 d.p.i. mice. **f,** Heatmap showing the differential expression of Mac2 specific genes identified by Remmerie *et al.*^14^. Data from 1,200 naïve cells and 6,152 cells from 42 d.p.i. mice, after QC filters.

Functionally, in infected mice the ResKCs cluster expressed several genes associated with KC roles, including lipid and iron metabolism^22^ (Fig. 5a), high levels of *Cxcl13* and *Ccl24*, which are expressed by steady state TRMs in different tissues^22,41^, and *Il10*, that has been associated with *Leishmania* persistence^42–44^. However, even though most resKCs maintained their putative function, they also appeared responsive to the Th1 environment of the infected liver^45,46^, as evidenced by expression of *Cxcl10*, *Cxcl9* and *Il1a* (Fig. 5b,g). MoKCs and Mac1 clusters showed a chemokine/cytokine profile that resembled more the resKCs than the macrophages in other clusters, but did not express most genes associated with lipid and iron metabolism (Fig. 5b). Transitioning monocytes1 showed the highest expression of several pro-inflammatory chemokines and cytokines, including *Tnf*, *Il1b*, *Cxcl9*, *Cxcl10*, *Ccl4* and *Il1a* (Fig. 5b). By confocal microscopy, iNOS expression peaked at 42 d.p.i. (Fig.5c-d and Extended Data Fig. 6a) and was restricted to resKC-derived granuloma macrophages and CLEC4F^-^TIM-4^-^ momacs within 42 d.p.i. granulomas (Fig. 5e-f). Thus, the *Nos2* annotated UMAPs will have identified macrophages that were present in granulomas, and included cells from Transitioning monocytes1 and 2, and Mac2 clusters (Fig. 5g and Extended Data Fig. 5b). The highest expression of *Cxcl10* and *Tnf* were also detected in the same *Nos2* expressing clusters (Fig. 5g), reinforcing the pro-inflammatory nature of the granuloma. By contrast, *Ccl24* and *Il10* were detected mostly in cells from ResKCs and MoKCs clusters (Fig. 5g). Altogether, Transitioning monocytes1 and 2 and Mac2 clusters expressed iNOS and high levels of pro-inflammatory cytokines and chemokines inside the granulomas, while both ResKCs and MoKCs showed a more homeostatic/regulatory profile, expressing the highest levels of IL-10. Thus, there is considerable functional heterogeneity among KCs and recruited macrophages that likely reflects their ontogeny, length of time and spatial distribution within the infected liver.

**Figure 5:**
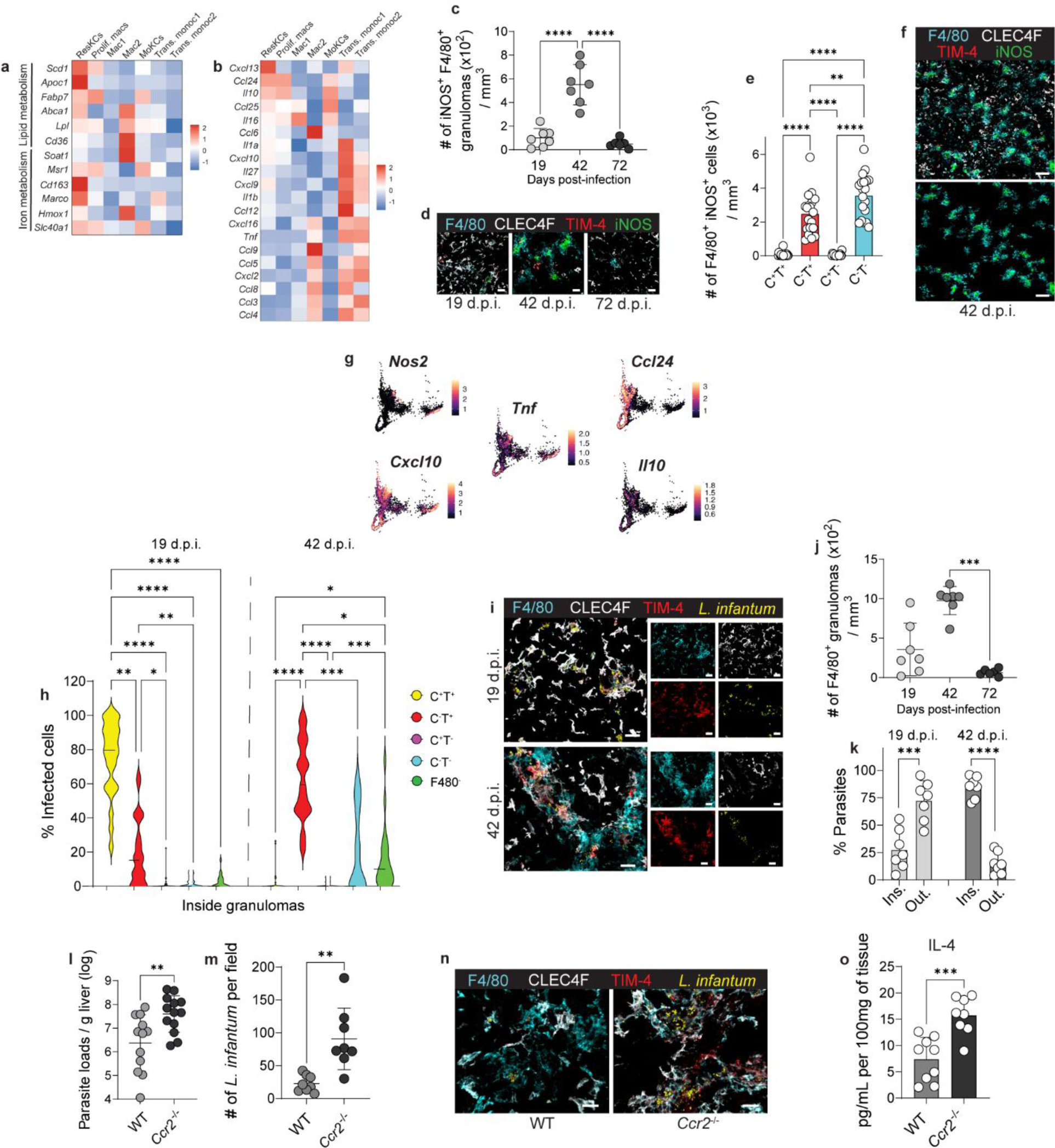
Functional heterogeneity of macrophage subsets and the contribution of of monocyte-derived macrophages to *L. infantum* control. **a-b**, Heatmaps showing the average expression levels of lipid and iron metabolism genes (**a**), and chemokines and cytokines (**b**) at 42 d.p.i. **c,** Number of iNOS^+^F4/80^+^ granulomas at different times post-infection in WT mice, quantified from immunofluorescence images. **d**, Representative images showing F4/80 (cyan), CLEC4F (white), TIM-4 (red), and iNOS (green) in WT 19-, 42-, and 72-day infected livers. Scale bars, 30μm. **e**, Number of F4/80^+^iNOS^+^ macrophages, based on CLEC4F and TIM-4 expression in infected WT livers at 42 d.p.i., and quantified from immunofluorescence images. Data pooled from 5 independent experiments (n=19). **f**, Representative images showing F4/80 (cyan), CLEC4F (white), TIM-4 (red), and iNOS (green) above, and only F4/80^+^iNOS^+^ granulomas below, in a WT liver at 42 d.p.i. **g**, UMAP plots showing the single-cell expression of *Nos2*, *Cxcl10*, *Tnf*, *Ccl24,* and *Il10* in CD45.2^+^F4/80^+^CD11b^int^CD64^+^ cells from 42-day infected mice. **h**, Frequency of infected cells found inside F4/80^+^ granulomas, at 19 and 42 d.p.i., and obtained from immunofluorescence images. **i**, Representative images of F4/80^+^ macrophages in WT 19- and 42-day infected livers, stained with anti-F4/80 (cyan), anti-CLEC4F (white), anti-TIM-4 (red), and anti-*L. infantum* (yellow). Scale bars, 30 μm. Data pooled from 2 independent experiments for each time point (n=8 for 19- and 42 d.p.i., n=7 for 72 d.p.i.), and from images of 2-3 regions from each liver containing granulomas (ROIs=24 for 19 d.p.i., ROIs=23 for 42 d.p.i., ROIs=16 for 72 d.p.i.). **j**, Number of F4/80^+^ granulomas quantified from immunofluorescence images. Data pooled from 2 independent experiments (n=7 for 19- and 42 d.p.i. and n=6 for 72 d.p.i.). **k**, Distribution of parasites inside and outside F4/80^+^ granulomas at 19 and 42 d.p.i., quantified from immunofluorescence images. Data pooled from 2 independent experiments (n=7). **l**, Parasite loads in WT and *Ccr2*^-/-^ mice at 42 d.p.i. Data pooled from 3 independent experiments (n=12 for WT and n=13 for *Ccr2*^-/-^). **m**, Number of *L. infantum* amastigotes per field in 42 day-infected WT and *Ccr2*^-/-^ mice, quantified from confocal microscopy images. Data pooled from 2 independent experiments (n=8). **n**, Representative images showing F4/80 (cyan), CLEC4F (white), TIM-4 (red), and *L. infantum* (yellow) in WT and *Ccr2*^-/-^ 42-day infected livers. Scale bars, 20μm. **o**, IL-4 levels measured by Luminex in liver homogenates from WT and *Ccr2*^-/-^ mice at 42 d.p.i. Data pooled from 2 independent experiments (n=9 for WT and n=8 for *Ccr2*^-/-^).Values in **c**,**e**, **h**, **j-k**, **l-m** and **o** represent the mean ± SD. *, p < 0.05, **, p < 0.01, ***, p < 0.001, ****, p < 0.0001. Data shown in heatmaps **a-b** were downsampled to a maximum of 500 cells per cluster for a more even cell type representation.

We also used confocal microscopy to address which macrophage populations were infected. Intra-granuloma *L. donovani* amastigotes were previously found within KCs in infected livers (8-28 d.p.i.), which were marked as a resident population by their uptake of colloidal carbon or fluorescent nanobeads prior to infection^20,24^. Here, *L. infantum* was similarly found mainly inside CLEC4F^+^TIM-4^+^resKCs at 19 d.p.i. (Figs.5h-i and Extended Data Fig. 6a). At the peak of granuloma formation at 42 d.p.i. (Fig. 5j), amastigotes were mostly found in resKC-derived macrophages inside granulomas and not in the momacs (Fig. 5h-i,k and Extended Data Fig. 6f), suggesting that the resKCs that were parasitized at earlier time points remained the main source of infected cells in the granulomas observed outside of the sinusoids. This raises the question as to what role the uninfected, recruited cells are playing in the development of the anti-parasitic response in the liver. Deficient monocyte recruitment during *L. donovani* infection has been previously shown to result in disorganized granulomas and increased parasite burdens^47,48,49^. We also found that *Ccr2*^-/-^ mice had 16.8-fold more liver parasites than WT mice at 42 d.p.i. (Fig. 5l). Quantification by confocal microscopy, showed that resKCs and resKC-derived macrophages of the *Ccr2*^-/-^ mice contained 4-fold more parasites than WT mice (Figs. 5m-n and Extended Data Fig. 6c). This susceptibility was associated with granulomas containing only resKCs and resKC-derived macrophages and not momacs. In the absence of recruited macrophages, CLEC4F^+^TIM-4^+^resKCs and resKC-derived macrophages from *Ccr2^-/-^* mice were activated to express iNOS (Extended Data Figs.6d-e) but were still not able to contain the infection as effectively as the WT mice. While IFN-γ, TNF and IL-1β (Extended Data Fig. 6f) cytokine levels were not affected, infected *Ccr2*^-/-^ mice showed increased levels of hepatic IL-4 (Fig. 5o). Increased IL-4, along with the absence of pro-inflammatory momacs, could explain why the resKCs and resKC-derived granuloma macrophages were compromised in their ability to control the liver parasite burdens in *Ccr2*^-/-^ mice. The contribution of diffusible NO and activating cytokines from uninfected momacs within the granuloma may be required for an optimal concentration of NO to be reached, especially as it is known that individual cells expressing iNOS are not able to control *Leishmania* growth^50^.

In summary, by using CLEC4F and TIM-4 staining, confocal microscopy, fate mapping reporter mice, and scRNA-seq analysis, we have provided direct evidence for a remarkable liver macrophage heterogeneity during experimental VL, informing their ontogeny, spatial distribution, and function in granuloma maturation and immunity. VL results in two main hepatic compartments populated by distinct macrophage subsets. The sinusoidal compartment is composed of resKCs and moKCs that maintain a homeostatic/regulatory phenotype. The mature granuloma compartment is found outside of the sinusoids and contains infected iNOS^+^ CLEC4F^-^resKC-derived macrophages, and uninfected, pro-inflammatory monocyte-derived macrophages. The remodeling of the extra-sinusoidal space by granulomas constituted by migrating, reprogrammed resKC-derived macrophages and recruited macrophages is closely linked to the evolution of hepatic immunity in VL.

## ACKNOWLEDGMENTS

We thank O. Schwartz and M. Smelkinson (Biological Imaging Section) for help with microscope setup and image acquisition and analysis; C. Eigsti, T. Moyer and I. Douagi (Flow Cytometry Section) for help with the cell sorting; P. Duncker (Cytek Biosciences) for help with the flow cytometry panel design; and P’ng Loke for critical review of the manuscript.

This work was supported in part by the Intramural Research Program of the NIAID, National Institutes of Health.

## AUTHOR CONTRIBUTIONS

Conceptualization, methodology, and validation, G.P., A.P. E.P.A., P.H.G.G, S.G., S.H.L. and D.L.S.; Investigation, G.P., A.P., E.P.A., P.H.G.G, O.K.; Software, T.R.F., J.K., S. R. P.; Data curation, T.R.F.; Formal analysis, G.P, T.R.F and J.K.; Resources, D.L.S.; Writing - original draft, visualization, supervision, and project administration G.P. and D.L.S.; Writing – Review and Editing, G.P. and D.L.S.; Funding acquisition, D.L.S.

## METHODS

### Mice

All the mice employed in our study were female, 6-8 weeks old, and maintained at the NIAID animal care facility under specific pathogen-free conditions with water and food *ad libitum*. The experiments were approved by the NIAID Animal Care and Use Committee (protocol number LPD 68E), and all aspects of the use of animals in this work were monitored for compliance with The Animal Welfare Act, the PHS Policy, the U.S. Government Principles for the Utilization and Care of Vertebrate Animals Used in Testing, Research, and Training, and the NIH Guide for the Care and Use of Laboratory Animals. The following mice were used: C57BL/6NTac (Taconic), *Ccr2*^-/-^ Jackson line #004999 (Taconic #8456)^51^, *Caspase1*^-/-^ (Taconic #8460)^52^, *Clec4f-Cre-tdTomato* (JAX stock #033296)^12^, Ai6(RCL-ZsGreen) (JAX stock #007906), *R26R-Confetti* (JAX stock #013731)^53^, *Bach1*^-/-54^, *Mlkl*^-/-^ (JAX stock #037116)^55^. #8456 and #8460 were obtained through contract between the NIAID and Taconic.

### Parasites

*Leishmania infantum* (MHOM/ES/92/LLM-320; isoenzyme typed MON-1) was cultured in M199 medium supplemented with 20% fetal bovine serum (Gemini Bio-Products), 100 U/mL penicillin/100 μg/mL streptomycin (Gibco), 2 mM L-glutamine (Gibco), 40 mM HEPEs (Gibco), 0,1 mM adenine (Sigma) in 50 mM HEPEs, 5 mg/mL hemin (Sigma) in 50% triethanolamine (Sigma), and 1 mg/mL 6-biotin (Sigma) at 26°C. For mice infection, metacyclic promastigotes were purified from stationary phase cultures by Ficoll (Sigma) density gradient (8% and 20%) centrifugation adapted from Spath and Beverley^56^, and centrifuged at 500*xg* for 10 minutes, at 25°C. Mice were then intravenously inoculated with 3×10^6^ metacyclic parasites in 100μl 1x PBS.

### Leukocytes isolation and purification from the liver

Mice were euthanized in a CO_2_ chamber and immediately perfused with 20 mL sterile 1x PBS (Gibco). The gallbladder was removed, and the livers were weighed and maintained in 1x PBS at 4°C until processed. Livers were place in 3mL of a solution containing 0.5% collagenase IV (Worthington) and 0.5 mg/mL DNAse (Sigma) in RPMI (Gibco), for 30 minutes at 37°C. The digested livers were processed in a 70μm cell strainer with 1x PBS. For leukocytes purification, the cells were centrifuged in 34% Percoll (GE Healthcare) solution in 1x PBS, 500*xg*, for 20 minutes at 25°C. Red blood cell lysis was performed with ACK lysis solution (Lonza) and cells washed once with 1x PBS (Gibco).

### Immunolabeling for flow cytometry

All cells were stained with LIVE/DEAD Fixable Blue Dead Cell Stain Kit (Thermo Fisher Scientific), diluted 1:1000 in 1x PBS, for 20 minutes at 4°C, in the dark. Nonspecific labelling was blocked with 1:100 TruStain FcX™ (anti-mouse CD16/32) antibody (Biolegend), and the surface markers employed were: F4/80 (BM8, Biolegend) 1:200, CD11b (M1/70, Biolegend) 1:800, CD11c (N418, Biolegend) 1:200, Ly6C (HK1.4, Biolegend) 1:400, Ly6G (1A8, Biolegend) 1:200, Siglec F (S17007L, Biolegend) 1:200, MHCII (M5/114.15.2, Biolegend) 1:800, CLEC4F (3E3F9, Biolegend) 1:50, CD64 (X54-5/7.1, Biolegend) 1:200, TIM-4 (RMT4-54, eBioscience^TM^) 1:200, diluted in FACS buffer (0.5% FBS + 1mM EDTA in 1x PBS). The samples were washed once with FACS buffer, fixed and permeabilized with BD Cytofix/Cytoperm™ (BD Biosciences) for 30 minutes at 4°C in the dark, and washed once with BD Perm/Wash™ buffer. Intracellular antibodies, iNOS (CXNFT, eBioscience^TM^) 1:800, and Ki-67 (SolA15, eBioscience^TM^) 1:160, were diluted in BD Perm/Wash™ buffer and incubated for 30 minutes at 4°C in the dark, then washed twice with BD Perm/Wash™ buffer. Cells were resuspended in FACS buffer and AccuCheck Counting Beads (Thermo Fisher Scientific). Labelled leukocytes were acquired using the Cytek^®^ Aurora cytometer (Cytek^®^ Biosciences) and the analysis performed with SpectroFlo^®^ software (Cytek^®^ Biosciences, version 3.0.1) and FlowJo^TM^ software (Treestar^®^, version 10.8.1).

### Quantification of parasite loads

The livers were perfused with 20 mL 1x PBS, and leukocytes isolated as previously described here, and resuspended in 1.5-2.5 mL of 1x PBS. Parasite loads were quantified similarly to Lee et al., 2020^57^. Briefly, a volume of 30-60μL was used for 2-fold serial dilutions in a flat bottom 96-well plate containing 150μL of M199/S per well and kept at 26°C. After 7-10 days the presence of parasites in the wells was evaluated, and the parasite loads per gram of tissue was calculated according to the highest dilution in which *L. infantum* was detected.

### Parabiosis

CD45.1^+^ and CD45.2^+^ mice with similar weights were cohoused for 2 weeks and surgically connected as described^58^. Three weeks post-surgery both mice from the pair were infected as described above and euthanized 42 days post-infection for leukocyte isolation and flow cytometry.

### Intravital microscopy

All intravital imaging experiments were approved by and performed in accordance with the Animal Care and Use Committee of the National Institute of Allergy and Infectious Diseases. Intravital imaging of the liver was performed as previously described ^59^. Briefly, images were acquired using a Leica DIVE (Deep In Vivo Explorer) inverted microscope (Leica Microsystems) equipped with full range of visible light lasers (Spectra Physics) and 37°C environmental Chamber (NIH Division of Scientific Equipment and Instrumentation Services). Anesthesia was provided using SurgiVet vaporizer and a nose-cone mask (Braintree Scientific). Infected *Clec4f-Cre-ZsGreen* or *Clec4f-Cre-Confetti* mice were anesthetized with 2% isoflurane delivered into induction chamber (Braintree Scientific) and maintained at 1.75% during imaging. The imaging was performed in regular confocal mode using long-working distance objective (L HC FLUOTAR 25x, NA=0.95, Leica Microsystems). To visualize blood vessels in the liver, 0.025 mg of anti-CD31 AF647 (clone MEC13.3, Biolegend) was injected intravenously. The liver was surgically accessed, restricted using custom-built tissue holder, and immersed in pre-warmed lubricating jelly. Static images were tiled, and a merged mosaic was constructed using Leica tile scanning software. For image quantification, chosen cells were rendered as 3D surfaces objects in Imaris software (Andor, version 9.7.2). Channel intensity statistics were exported directly from Imaris as .ims files using python parser as csv datasets and imported into FlowJo^TM^ (Treestar®, version 10.8.1) for gating and analysis. Images were analyzed with Imaris software (Andor, version 9.7.2). Parser was written in python and source code is available at https://github.com/srperera/nih_parsers/tree/master/surface_parser.

### Anti-Leishmania antibody purification from serum

To visualize *L. infantum* amastigotes by confocal microscopy, we used pooled serum samples from kala-azar cured patients^60^, centrifuged at 10,000*xg* for 10 minutes at 4°C, and IgG purified using the Pierce™ Protein A IgG Purification Kit (Thermo Fisher Scientific) following the manufacturer’s instructions. Eluted antibodies were pooled and dialyzed in 1x PBS 3x for 2 hours and 1x overnight at 4°C. Antibodies were concentrated using the Amicon^®^ Ultra-4 50K (Merck Millipore) at maximum speed for 15 minutes, lyophilized, and stored at −80°C until use. Antibodies were resuspended in 1x PBS and conjugated with ReadiLink™ xtra Rapid iFluor^®^ 488 Antibody Labeling Kit (AAT Bioquest) and used for staining fixed/frozen liver sections.

### Confocal microscopy

We adapted the fixation and staining steps of the protocol described by Radtke *et al.*^61^. Briefly, livers were perfused in 1x PBS and a piece of the larger lobule was fixed with BD Cytofix/Cytoperm™ (BD Biosciences) diluted 1:4 in 1x PBS, for 24h at 4°C. The tissues were soaked in 30% sucrose (Sigma) solution overnight at 4°C, placed on a Histomold (Fisher Healthcare) and filled with Tissue-Tek^®^ O.C.T. Compound (Sakura Finetek). The histomold was frozen on dry ice and stored at −80°C. OCT-embedded tissues were sectioned in a cryostat (Leica Biosystems), and 10μm sections were prepared on VWR® Superfrost® Plus Micro slides (VWR Corporate Headquarters) and kept frozen at −80°C until use. For immunostaining, tissue sections were rehydrated with 1x PBS for 5 minutes at RT, and blocked with block buffer containing 1% TruStain FcX™ (Biolegend) and 0.3% Triton X-100 (Calbiochem^®^) in 1% BSA (Thermo Fisher Scientific) for 1 hour at 37°C. The samples were then incubated with the following antibodies: F4/80 (BM8, eBioscience™) 1:100, CLEC4F (3E3F9, Biolegend) 1:100, TIM-4 (RMT4-54, Biolegend) 1:200, iNOS (CXNFT, eBioscience™) 1:200, cleaved caspase 3 (5A1E, Cell Signaling) 1:100, and/or anti-*L. infantum* (purified and conjugated in house) 1:200, in a 1% BSA solution containing 0.3% Triton X-100 and 1:5000 Hoechst 33342 (Thermo Fisher Scientific) for 1h at 37°C. The tissue sections were then washed 5 times for 3 minutes with 1x PBS, and slides were mounted with the Prolong^TM^ Diamond Antifade Mountant (Thermo Fisher Scientific), following the manufacturer’s instructions. Tiled images of 3×4 for naïve and 4×4 tiles for infected mice were acquired using the Leica SP8 WLL FLIM confocal microscope. Tiles were merged using the LAS X Navigator software (LAS X 3.5.5.19976) and analyzed with Imaris software (Andor, version 9.7.2). Image quantification was performed as described above.

### Lipid peroxidation assay and intracellular GSH measurement

Lipid peroxidation in F4/80^hi^CD11b^int^CD64^+^TIM-4^+/-^ and F4/80^low/-^CD11b^+^ cells was assessed using BODIPY™ 665/676 (Lipid Peroxidation Sensor) (Thermo Fisher Scientific). Briefly, cells were isolated as described above and incubated with 10μM Bodipy in 1x PBS for 30 minutes at 37°C. Cells were washed once with 1x PBS and extracellular staining was performed as described above. Glutathione (GSH) levels were evaluated in liver homogenates. Briefly, livers were perfused with 20 mL 1x PBS and a piece of the larger lobule was homogenized using glass beads in 1x PBS, and centrifuged at maximum speed at 4°C for 10 minutes, to remove tissue matrix and cell debris. Supernatants were stored at −80°C until use. Reduced glutathione levels were measured using the Glutathione Assay Kit (Cayman, USA, 703002) following the manufacturer’s instructions. Total protein was measured by Pierce Protein Assay (Thermo Fisher Scientific), according to the manufacturer’s instructions, and values used to normalize GSH levels based on total protein content.

### Single-cell RNA sequencing

Liver leukocytes were isolated as described above, and tissue digestion/Percoll solution and staining buffers were supplemented with 100U/mL SUPERase•In™ RNase Inhibitor (Thermo Fisher Scientific). Live, single CD45.2^+^ F4/80^hi^ CD11^int^ CD64^+^ cells were sorted using the Cytek^®^ Aurora CS sorter and 3,000 cells were used for RNA extraction and library preparation using the Chromium Next GEM Single Cell 3’ Kit v3.1 (PN-1000269, 10x Genomics) according to the manufacturer’s instructions. Briefly, single cells were isolated in aqueous droplets containing beads coated with barcoded oligo-dT in a water-in-oil emulsion using a Chromium controller (10x Genomics). Transcripts from lysed cells were reversed transcribed into cDNA and processed by amplification, fragmentation, dual-adapter ligation and size selection. Libraries were then analyzed on a D5000 ScreenTape using a TapeStation (Agilent) and sequenced on an Illumina HiSeqX Ten by Psomagen Inc.

Sequence reads were demultiplexed into FASTQ files from raw base calls using the *mkfastq* module in Cell Ranger v.5.0.0 (10x Genomics). Raw FASTQ files were quality checked using FastQC and filtered for reads with average quality of 20 and minimum length of 20bp using fastp v.0.20.^62^. Cell Ranger v.7.1.0 *count* was used to align filtered reads to the mouse reference transcriptome mm10 (GENCODE vM23/Ensemble 98) and to quantify the number of different unique molecular identifiers (UMIs) for each gene. The fraction of reads containing a valid barcode assigned to a cell and confidently mapped to the transcriptome varied between 45.3-60.6% in naive samples and 86.7%-89.3% in infected samples. The estimated number of cells detected in reach replicate was between 1,166 and 4,804 cells, for a total of 11,834 cells used as input loaded into RStudio v.2023.03.0 for downstream processing and analysis using Seurat v.4.3.0^63^.

Cells were removed from the analysis if either the mitochondrial gene expression represented > 10% of total expression, displayed expression < 1000 different genes, > 6000 different genes or < 5000 transcript molecules. Genes detected in fewer than 2 cells were also removed. UMI counts were normalized by library read depth, log-transformed, centered and Z-scored using functions *NormalizeData* and *ScaleData*. Highly variable genes were identified using *FindVariableFeatures* (2000 features, “vst”method). Data integration was performed by selecting genes that are variable in most samples with *SelectIntegrationFeatures* and identifying cells to be used as “anchors” with *FindIntegrationAnchors*, to integrate the different samples using the *IntegrateData* function. Dimensionality reduction was performed on scaled integrated data using *RunPCA* and *RunUMAP* and clustering with *FindNeighbors* and *FindClusters* with 15 principal components and a resolution of 0.2. To be consistent with annotation from the literature on KCs, we mapped our data onto a reference scRNA-seq dataset from Remmerie *et al.*^14^ using the *MapQuery* function. Transcript markers for each cell cluster were identified using the *FindAllMarkers* function and evaluating the average expression and the percent expressed of each gene in a cluster compared to others. Markers were selected if single-cell log2FC expression > 0.25 and statistically significant (Wilcoxon rank-sum test adjusted p-value < 0.05). Clusters were manually annotated according to their mapping to the reference dataset. KC signature score was calculated using the *AddModuleScore* iunction with the final list of 15 human-mouse conserved KC markers previously described {REF} (*Cd5l, Vsig4, Slc1a3, Cd163, Folr2, Timd4, Marco, Gfra2, Adrb1, Tmem26, Slc40a1, Hmox1, Slc16a9, Vcam1, Sucnr1*).

### Multiplex cytokine arrays

To evaluate the inflammatory mediators in the liver homogenates across the groups, approximately 45mg of liver were homogenized for 2 cycles of 20 seconds at 5000 rpm each using Precellys tubes with metal beads in 200μL of RIPA lysis buffer (Millipore) supplemented with protease inhibitor cocktail (Thermo Fisher Scientific). Following centrifugation at 8000*xg* for 10 minutes at 4°C, the supernatant was used to measure cytokines in the liver homogenate. The cytokines IL-4, IL-1b, IFN-γ, and TNF-α levels were measured using the commercial MILLIPLEX MAP mouse cytokine/chemokine magnetic bead panel kit (Millipore Sigma) according to manufacturer instructions, and the cytokines values in pg/mL were normalized by mg of liver tissue weight.

## STATISTICAL ANALYSES

Statistical analyses were performed with GraphPad Prism software version 10.0.0 (La Jolla). Data are expressed as mean ± standard deviation and were considered statistically significant when p < 0.05. In all analyses the normal distribution and the homogeneous variance were tested. When following normal distribution, unpaired t-student (two-tailed) or one-way ANOVA with Tukey’s multiple tests were applied. For the data with the non-normal distribution, unpaired Mann-Whitney (two-tailed) and Kruskall-Wallis tests were performed with Dunn’s post-test.

## DATA AND CODE AVAILABILITY

Raw data were deposited at NCBI SRA repository under the BioProject ID PRJNA994082. Seurat objects containing processed dataset are available on Zenodo: 10.5281/zenodo.10780584.

---

**Extended Data Fig.1:**
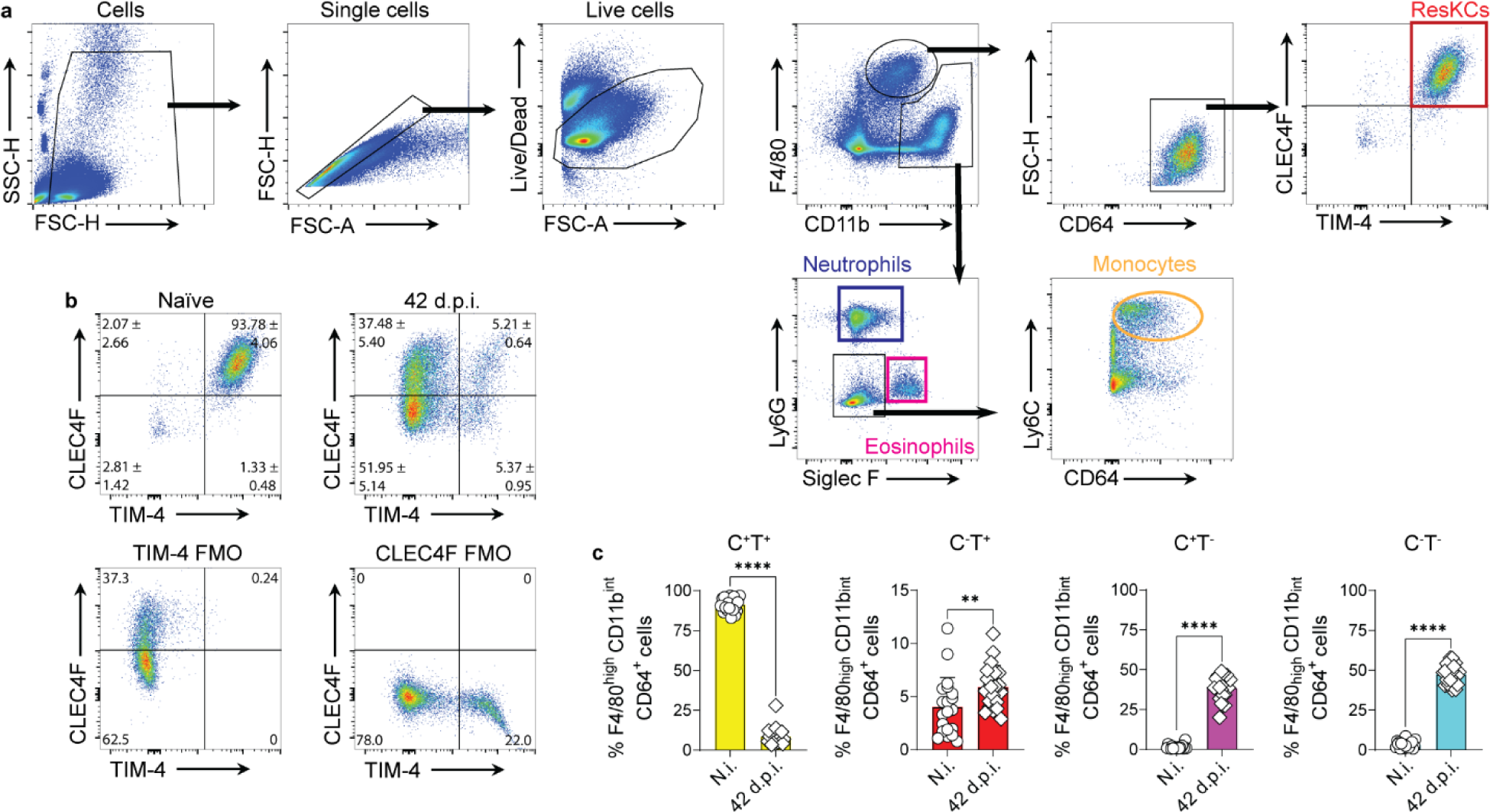
Myeloid cells gating strategy and macrophage heterogeneity at 42 days post-infection. **a**, Dot plots showing the gating strategy used to identify resKCs, (F4/80^hi^CD11bi^nt^CD64^+^CLEC4F^+^TIM-4^+^), neutrophils (CD11b^+^Ly6G^+^SiglecF^-^), eosinophils (CD11b^+^Ly6G^-^SiglecF^+^) and monocytes (CD11b^+^Ly6G^-^SiglecF^-^Ly6C^+^CD64^+^) in live, single cells isolated from naïve WT livers. **b**, Representative FACS plots showing CLEC4F and TIM-4 expression gated on F4/80^hi^CD11b^int^CD64^+^ macrophages in naïve and 42 day-infected mice, and fluorescence minus one (FMO) staining for CLEC4F and TIM-4. FMO controls contain mixed naïve and infected samples. **c**, Frequency of macrophage subsets in naïve and 42-day infected mice by flow cytometry. Numbers indicate mean ± SD percentage of cells in the gate. Data pooled from 5 independent experiments (n=20). Values in **c** represent the mean ± SD. **, p < 0.01, ****, p < 0.0001.

**Extended Data Fig.2:**
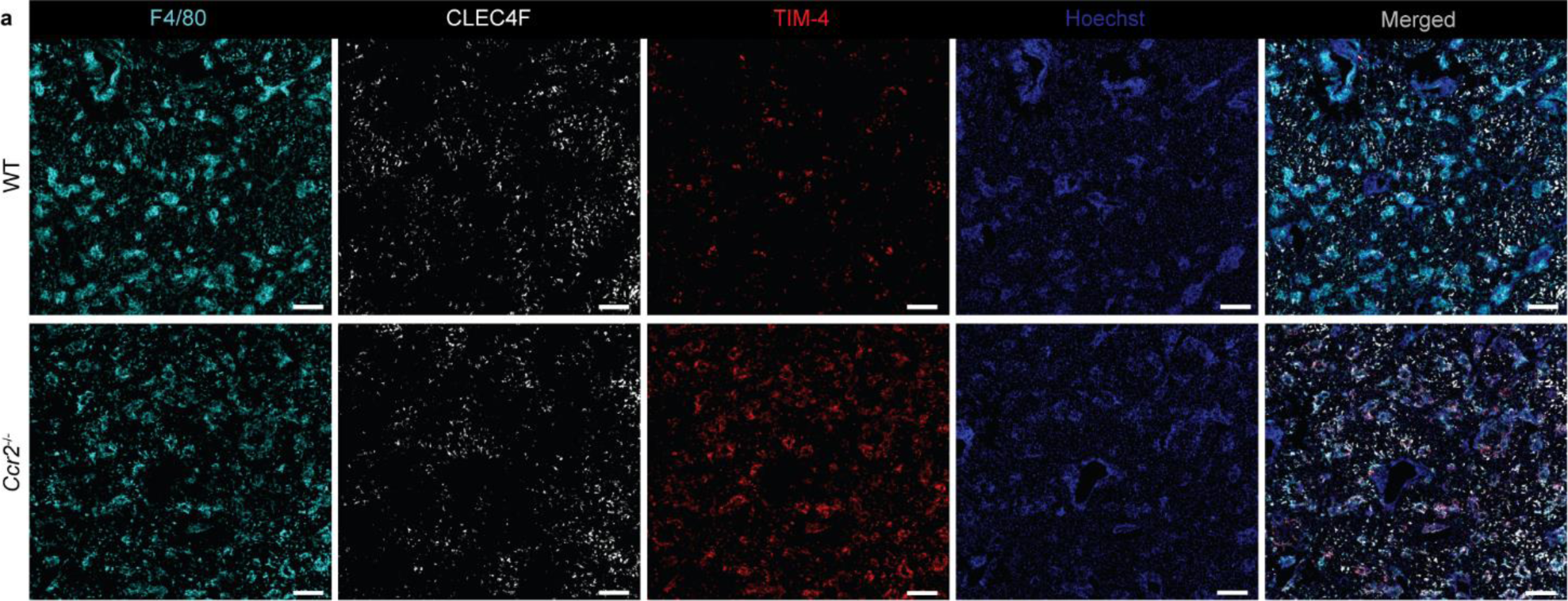
WT and *Ccr2*^-/-^ infected livers. **a**, Original representative immunofluorescence images of rendered images from **Fig**.**2i** showing WT and *Ccr2*^-/-^ 42 d.p.i. livers, and F4/80 (cyan), CLEC4F (white), TIM-4 (red), and Hoechst (blue). Scale bars, 200 μm.

**Extended Data Fig.3:**
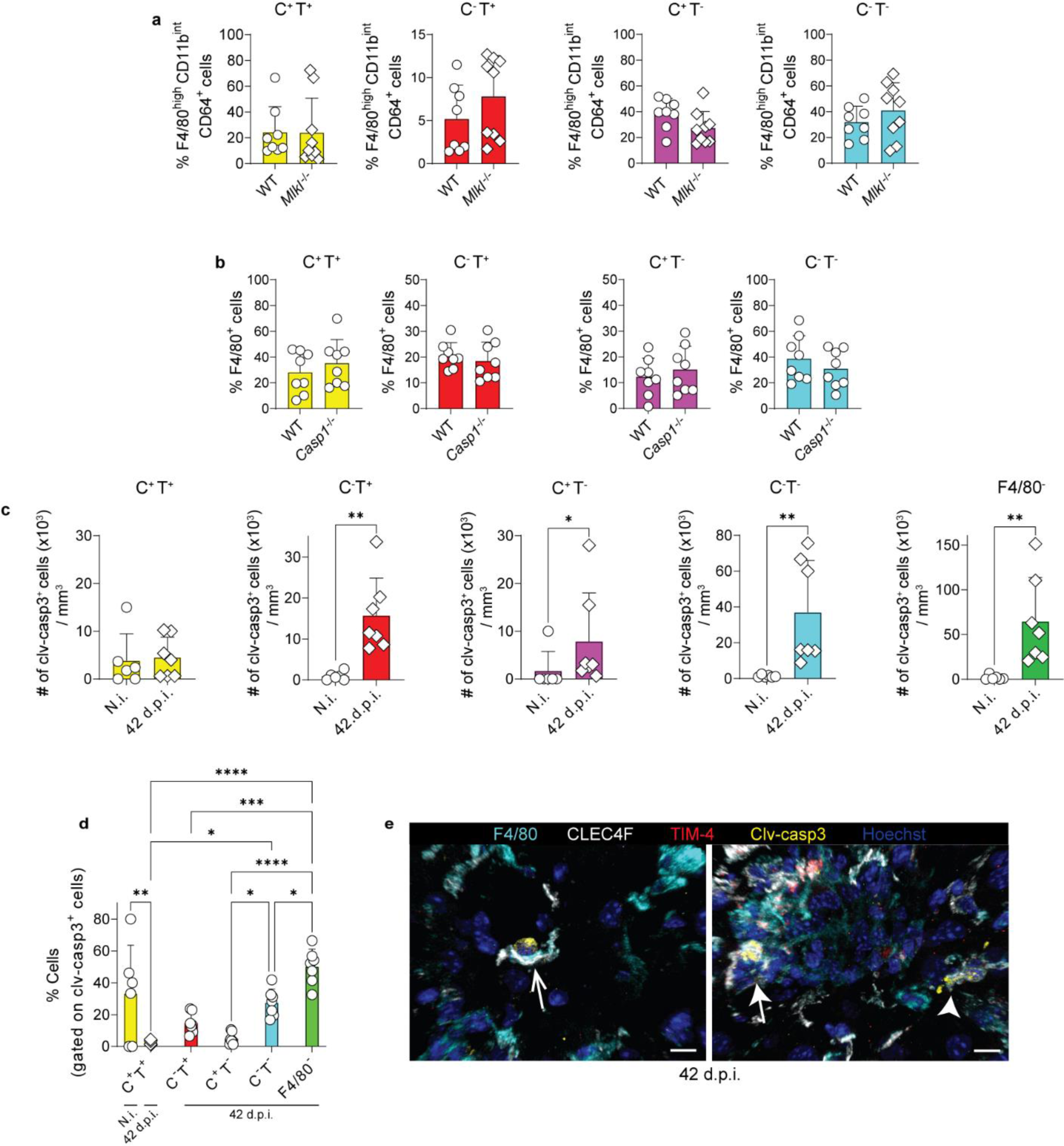
Cell death pathways in macrophages during VL. **a**, Frequency of cells in F4/80^hi^CD11b^int^CD64^+^ subsets in 42 d.p.i. WT and *Mlkl*^-/-^ mice, by flow cytometry. Data pooled from 2 independent experiments (n=8 for WT and n=9 for *Mlkl*^-/-^). **b**, Frequency of F4/80^+^ subsets in 42 d.p.i. WT and *Casp1*^-/-^ mice, quantified from immunofluorescence images (n=8). **c**, Number of cleaved caspase 3^+^ cells in each F4/80^+^ subset and in F4/80^-^ cells quantified from immunofluorescence images in naïve and 42-day infected WT mice. **d**, Frequency of each subset of F4/80^+^ and F4/80^-^ cells, gated on cleaved caspase 3^+^ cells, in naïve and 42-day infected WT mice. **e**, Representative immunofluorescence images showing apoptotic CLEC4F^+^TIM-4^-^ moKC (thin arrow), apoptotic CLEC4F^-^TIM-4^+^ resKC-derived macrophage (arrow), and apoptotic CLEC4F^+^TIM-4^+^ resKC (arrowhead) in 42-day infected mice. Scale bars, 8 and 10 μm, respectively. Data pooled from 2 independent experiments (n=6 for naïve and n=7 for 42 d.p.i.). Values in **a-d** represent mean ± SD. *, p < 0.05, **, p < 0.01, ***, p < 0.001, ****, p < 0.0001.

**Extended Data Fig.4:**
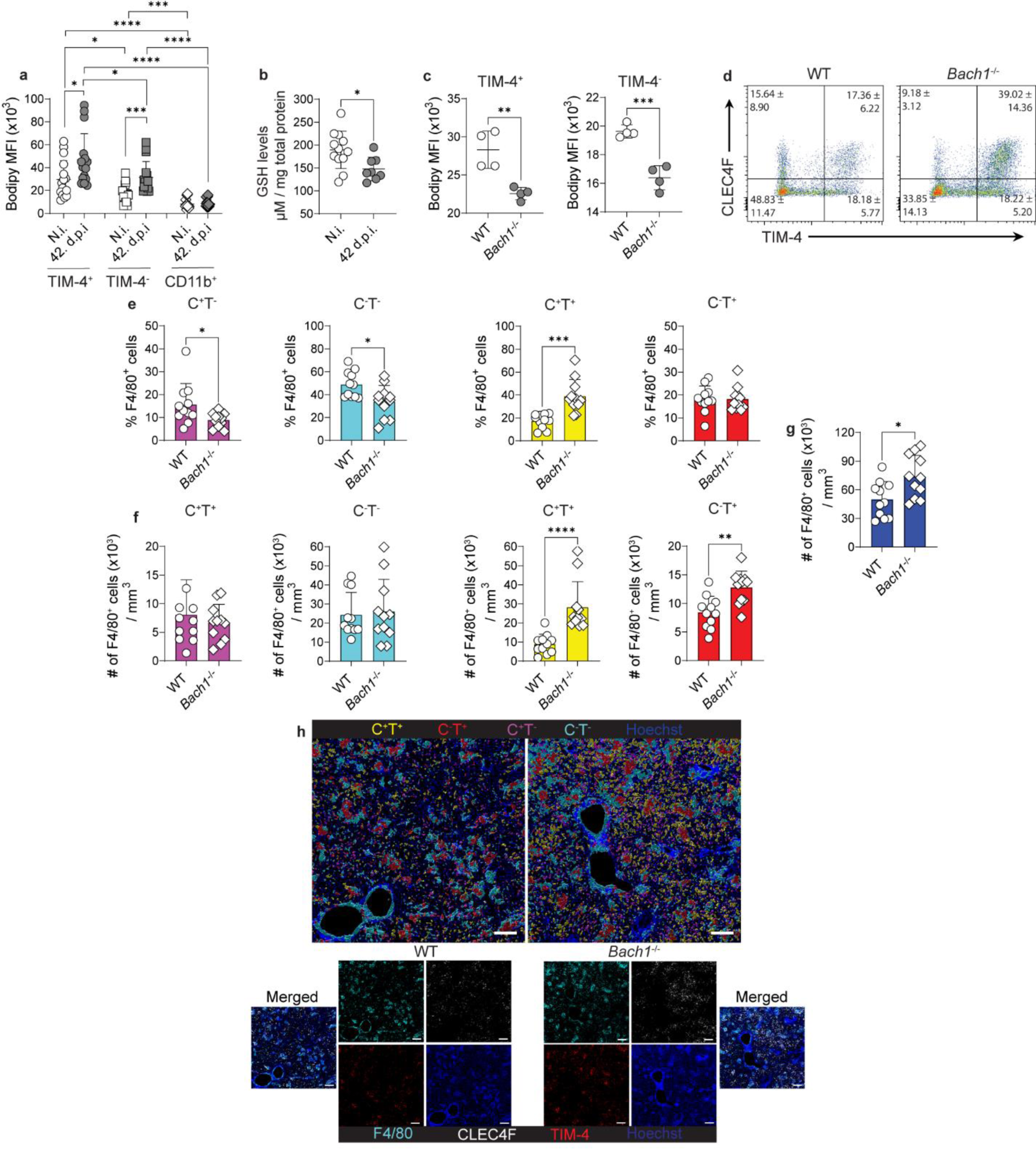
Ferroptosis contributes to reduction in resKC numbers during VL. **a**, Lipid peroxidation measured by bodipy median fluorescence intensity in TIM-4^+/-^ cells (gated on F4/80^hi^CD11b^int^CD64^+^ macrophages) and in CD11b^+^ cells (excluding F4/80^hi^ macrophages), in WT naïve and 42 d.p.i. by flow cytometry. Data pooled from 4 independent experiments (n=16). **b**, Intracellular GSH levels measured in whole tissue homogenates from naïve and 42 d.p.i. WT mice. Data pooled from 2-3 independent experiments (n=12 for naïve and n=8 for 42 d.p.i.). **c**, Lipid peroxidation in TIM-4^+^ and TIM-4^-^ cells from WT and *Bach1*^-/-^ mice at 42 d.p.i. Data representative of 2 independent experiments (n=4 each). **d**, Representative dot plots from immunofluorescence images showing the frequencies of macrophages based on CLEC4F and TIM-4 expression in WT and *Bach1*^-/-^ at 42 d.p.i. Numbers indicate mean ± SD percentage of cells in the gate. **e-f** Frequency (**e**) and number (**f**) of F4/80^+^ subsets in 42 d.p.i WT and *Bach1*^-/-^ mice, quantified from immunofluorescence images. **g**, Number of F4/80^+^ macrophages in WT and *Bach1*^-/-^ mice, at 42 d.p.i., quantified from immunofluorescence images. **h**, Representative rendered images of WT and *Bach1*^-/-^ 42 d.p.i. livers showing CLEC4F^+^TIM-4^+^resKCs (yellow), CLEC4F^-^TIM-4^+^resKC-derived macrophages (red), CLEC4F^+^TIM-4^-^moKCs (magenta) and CLEC4F^-^TIM-4^-^momacs (cyan), and original immunofluorescence images showing F4/80 (cyan), CLEC4F (white), TIM-4 (red) and Hoechst (blue). Scale bars, 200 μm. Data pooled from 3 independent experiments (n=11). Values from **a-c** and **e-g** represent mean ± standard deviation. *, p < 0.05, **, p < 0.01, ***, p < 0.001, ****, p < 0.0001.

**Extended Data Fig.5:**
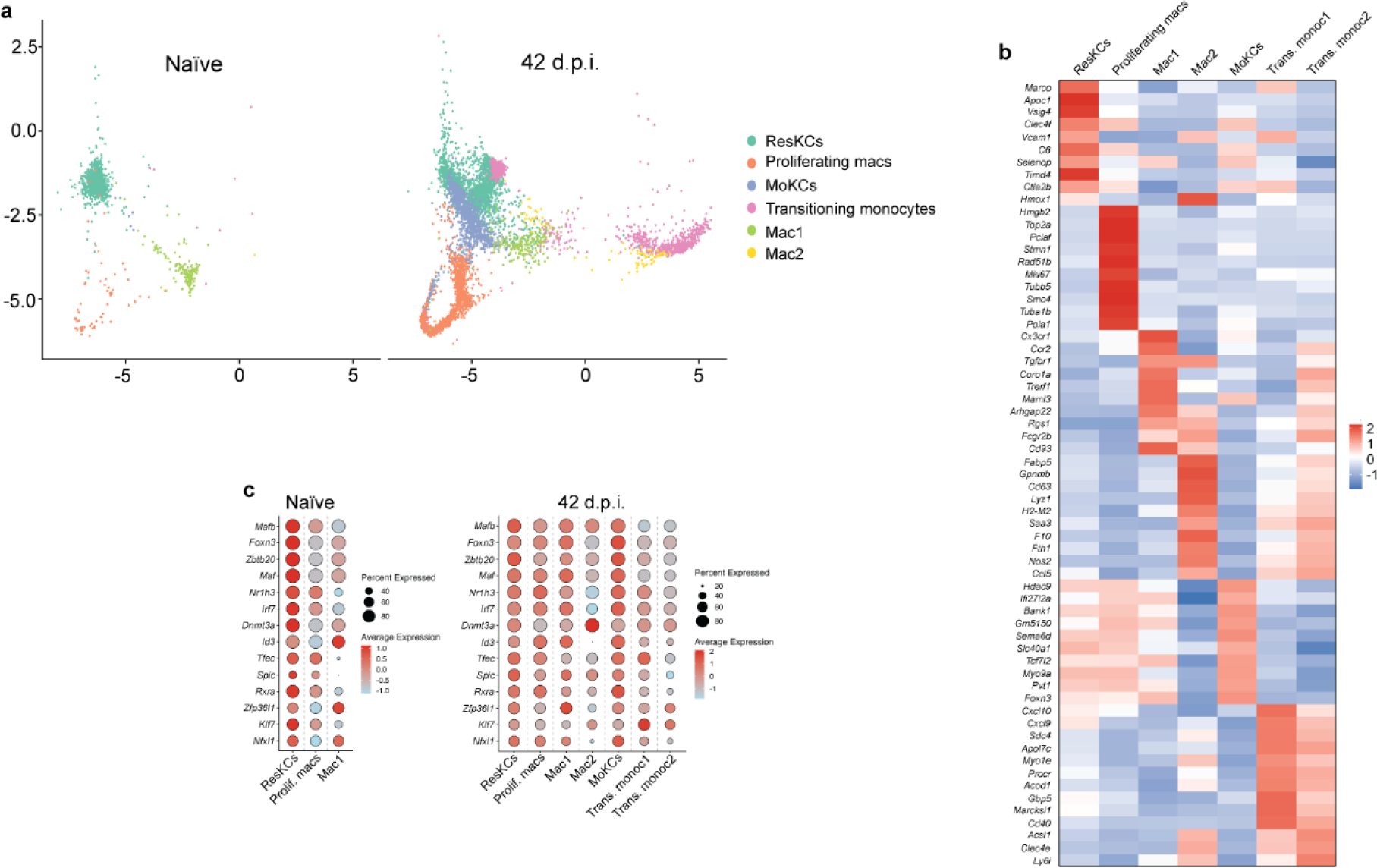
Clusters and differentially expressed genes identified by single-cell RNA sequencing. **a,** UMAP plot of scRNA-seq data of sorted live, single, CD45.2^+^F4/80^+^CD11b^int^CD64^+^ cells from uninfected and 42 d.p.i. mice, showing 3 clusters in naïve and 6 clusters in infected mice, as defined by Remmerie *et al.*^14^. **b**, Heatmap showing the average expression of the top 10 differentially expressed genes defining the clusters from sorted live, single, CD45.2^+^F4/80^+^CD11b^int^CD64^+^ cells of uninfected and 42 d.p.i. mice. **c**, Average gene expression and corresponding cell percent of KC-associated transcription factors within each of the clusters. Data from 1,200 naïve cells and 6,152 cells from 42 d.p.i. mice, after QC.

**Extended Data Fig.6:**
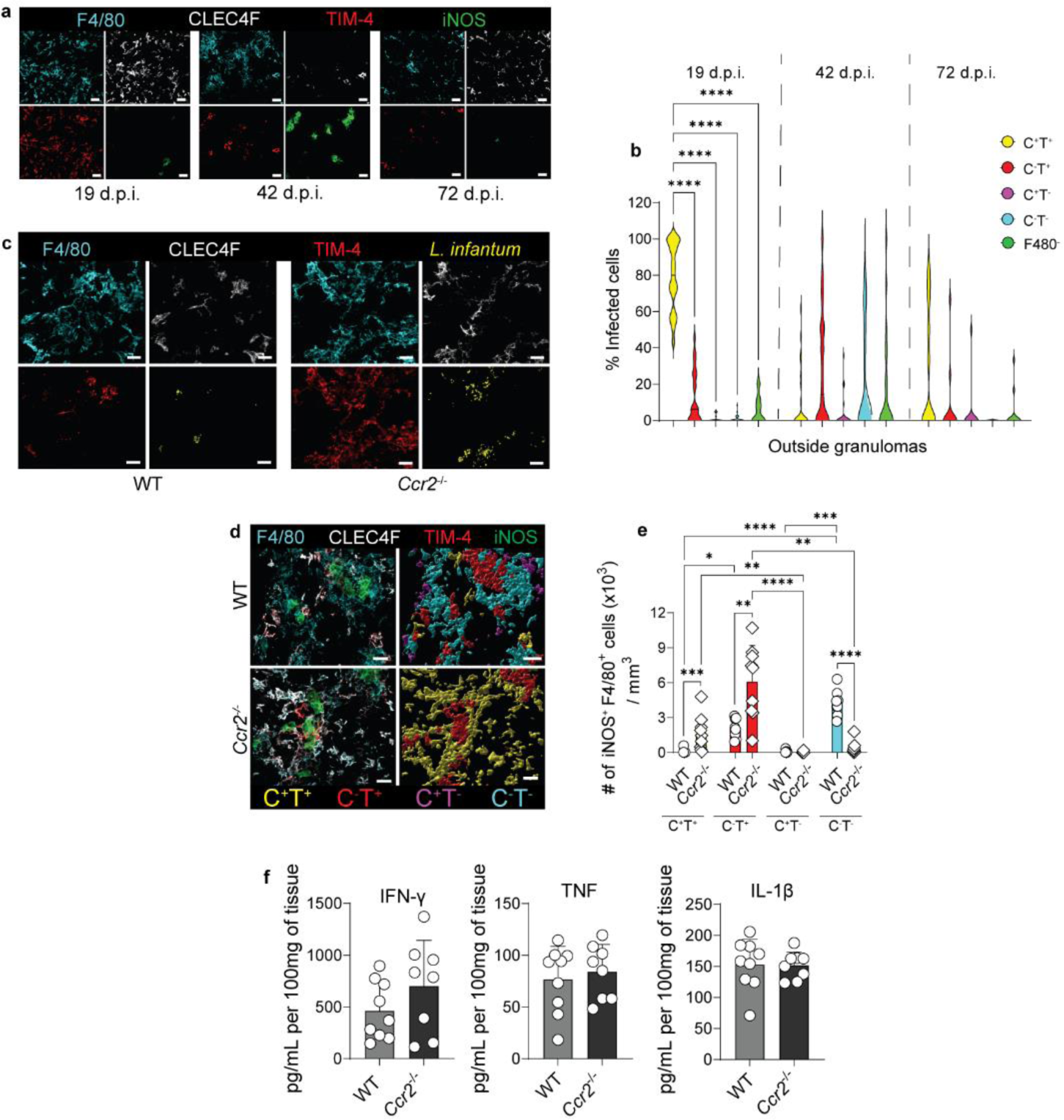
Activation and infection status of macrophage subsets. **a**, Individual images from **Fig.5d** showing F4/80 (cyan), CLEC4F (white), TIM-4 (red), and iNOS (green) in WT 19-, 42-, and 72-day infected livers. Scale bars, 30μm. **b**, Frequency of infected cells identified outside F4/80^+^ granulomas at 19 and 42 d.p.i., obtained from immunofluorescence images. Data pooled from 2 independent experiments for each time point (n=8 for 19- and 42 d.p.i., n=7 for 72 d.p.i.), obtained from images of 2-3 regions from each liver containing granulomas (ROIs=24 for 19 d.p.i., ROIs=23 for 42 d.p.i., ROIs=16 for 72 d.p.i.). **c**, Individual images from merged images in **Fig.5n** showing F4/80 (cyan), CLEC4F (white), TIM-4 (red), and *L. infantum* (yellow) in WT and *Ccr2*^-/-^ 42-day infected livers. Scale bars, 20μm. **d**, Representative and rendered images of F4/80^+^ granulomas and iNOS expression in WT and *Ccr2*^-/-^ 42-day infected mice, showing F4/80 (cyan), CLEC4F (white), TIM-4 (red), and iNOS (green) staining, and rendered CLEC4F^+^TIM-4^+^ (yellow), CLEC4F^-^TIM-4^+^ (red), CLEC4F^+^TIM-4^-^ (magenta), and CLEC4F^-^TIM-4^-^ (cyan) F4/80^+^ cells. Scale bars, 30μm. **e**, iNOS expression by different F4/80^+^ subsets in WT and *Ccr2*^-/-^ 42 d.p.i. mice, quantified from immunofluorescence images. **f**, IFN-γ, TNF and IL-1β levels measured by Luminex in liver homogenates from WT and *Ccr2*^-/-^ mice at 42 d.p.i. Data pooled from 2 independent experiments (n=9 for WT and n=8 for *Ccr2*^-/-^). Values in **b**, **e**-**f** represent the mean ± SD. *, p < 0.05, **, p < 0.01, ***, p < 0.001, ****, p < 0.0001.

